# Bcl-2 supports survival and metabolic fitness of quiescent tissue-resident ILC3

**DOI:** 10.1101/2023.02.27.528945

**Authors:** James I. King, Felipe Melo-Gonzalez, Bert Malengier-Devlies, Roser Tachó-Piñot, Marlene S Magalhaes, Suzanne H. Hodge, Xavier Romero Ros, Rebecca Gentek, Matthew R. Hepworth

## Abstract

Group 3 innate lymphoid cells (ILC3) are potent effector cells with critical roles in enforcing immunity, barrier integrity and tissue homeostasis along the gastrointestinal tract. ILC3 are considered to be primarily tissue-resident cells, seeding the gastrointestinal tract during embryonic stages. However, the mechanisms through which ILC3 are maintained within these tissues are poorly understood. Here, we report that ILC3 are minimally replenished from bone marrow precursors in healthy adult mice, persist in the tissue for extended periods of time in the gut, and display a quiescent phenotype. Strikingly, despite robustly producing cytokines, LTi-like ILC3 remain non-proliferative during enteric bacterial infection. Survival of LTi-like ILC3 was found to be dependent upon the balance of the metabolic activity required to drive effector function and anti-apoptotic programs. Notably, the pro-survival protein Bcl-2 was required for the survival of LTi-like ILC3 but was rendered partially dispensable if mitochondrial respiration was inhibited. Together we demonstrate LTi-like ILC3 are a quiescent-like population that persists independently of haematopoietic replenishment to survive within the tissue microenvironment.

## Introduction

The gastrointestinal tract represents a highly dynamic microenvironment in which the layered immune system acts to maintain tissue health and homeostatic function [1–3]. As a result, many intestinal immune cell populations turnover rapidly and are reliant on continued replenishment or supplementation from circulating bone marrow-derived progeny [4–6]. In contrast, the turnover of mature innate lymphoid cells (ILC) within tissue microenvironments remains incompletely defined, although the majority are thought to be primarily tissue-resident and non-circulatory, with the exception of natural killer (NK) cells and ILC1 [7,8]. In contrast, migration between the gut and organised intestinal associated lymphoid tissues can occur via the lymphatics, suggesting ILC may indeed leave the tissue and enter other organs in some contexts [9,10]. Group 3 innate lymphoid cells (ILC3), characterised by expression of the transcription factor retinoic acid-related orphan receptor gamma (RORγt), are amongst the first lymphocyte populations to establish residency in the gut and play important roles in driving lymphoid tissue organogenesis, and maintain intestinal barrier health throughout the life course via constitutive production of the cytokine interleukin (IL)-22 [11]. Recent advances have further defined the early ontological events that determine ILC3 seeding prior to birth and in the neonatal period [12–14], however the mechanisms intestine-resident ILC3 utilize to persist throughout adulthood, and the extent to which the bone marrow contributes to maintenance of this population remains poorly understood.

Intestinal ILC3 exhibit extensive heterogeneity and comprise at least two distinct subsets with significant differences in tissue localisation, transcriptional regulation, cell surface phenotype and biological functions [15]. Specifically, ILC3 can be split into two main subsets referred to as natural cytotoxicity receptor-expressing (NCR^+^) ILC3 and lymphoid-tissue inducer-like (LTi-like) ILC3, both of which mediate protective effector responses to extracellular microbes via the production of IL-22 [15]. Despite sharing overlapping effector cytokine profiles, NCR^+^ ILC3 and LTi-like ILC3 have the capacity to perform vastly divergent immune functions. LTi-like ILC3 are localized specifically within lymph nodes and intestinal-associated lymphoid structures [9,16–18] and possess a variety of regulatory functions that act to modulate adaptive immunity at mucosal barrier surfaces [19]. In contrast, NCR^+^ ILC3 are typically dispersed within the intestinal lamina propria where they establish residence following early life microbial colonisation events [20–22]. Moreover, unlike LTi-like ILC3, NCR^+^ ILC3 co-express the transcription factor T-bet which confers the capacity to secrete IFN-γ, and in inflammatory contexts NCR^+^ ILC3 can lose RORγt expression and conversion of NCR^+^ ILC3 to an ‘ex-ILC3’ or ILC1-like pro-inflammatory phenotype [23–25]. However these subset-specific differences in biology are often overlooked, and as a result ILC3 have been attributed seemingly contradictory roles in the context of intestinal immunity and inflammatory disease, provoking the need for a better understanding of the differences in ILC3 subset-specific biology in health and disease.

Here we show that intestinal ILC3 are long-lived tissue-resident cells that display relatively little hematopoietic replenishment under steady state conditions. We report LTi-like ILC3 exhibit a quiescent-like phenotype and rely on the anti-apoptotic protein Bcl-2, which facilitates survival in part by protecting cells against OXPHOS-mediated metabolic stress. Together these findings highlight LTi-like ILC3 as long-lived tissue-resident effector cells that are adapted for persistence within their tissue niche and further define the contrasting biology underpinning the immune functions of ILC3 subsets within mucosal barrier tissues - which may shed light on their roles in inflammatory diseases, such as IBD.

## Results

### Intestinal ILC3s are maintained independently of bone marrow replenishment

Previous studies have demonstrated that mature ILC do not typically recirculate in the blood [7], suggesting they may be maintained within tissues through mechanisms that remain unclear. To further define the turnover of ILC3 subsets within the intestine and associated lymphoid structures, we first utilized labelling of intestinal ILC with a previously described tamoxifen-inducible fate mapping model under the control of the *Id2* allele, a transcription factor common to all ILCs (*Id2*^CreERT2-RFP^) [26,27]. While the frequencies of RFP^+^ cells amongst both ILC3 subsets were observed to decrease slightly over a six week period, the vast majority of RFP^+^ cells were surprisingly maintained - suggestive of a relatively low turnover of ILC3 (Figure S1A–B). To address the bone marrow contribution towards the tissue-resident ILC compartment in adults, we next generated bone marrow chimeras in which the gastrointestinal tract was shielded to protect tissue-resident cells and analysed the contribution of congenic donor bone marrow (CD45.2^+^) to intestinal lymphocyte populations (CD45.1^+^) at least 10 weeks post-irradiation (Figure 1A). Notably, while other tissue-resident lymphocytes including CD4^+^ T cells exhibited approximately 30% chimerism over this time period, we found relatively low chimerism amongst both NCR^+^ ILC3 and LTi-like ILC3 across the intestinal tract and associated lymph nodes (Figure 1B-C). These findings suggest adult ILC3 populations exhibit a relatively reduced contribution from bone marrow haematopoiesis when compared to other lymphocyte populations at steady state. To determine whether bone marrow replenishment could occur following depletion of the tissue-resident ILC3 population, we next exposed mice to sub-lethal irradiation in the absence of intestinal shielding (Figure S1C). In this context bone marrow-mediated replenishment of the ILC3 compartment was significantly elevated (Figure S1D-E), indicating the bone marrow is capable of replenishing intestinal ILC3 following significant disruption of the cell population and/or tissue microenvironment, however this does not typically occur at a high rate in otherwise healthy. Together these data suggest intestinal ILC3 may be long-lived tissue-resident cells at homeostasis.

**Figure 1.**
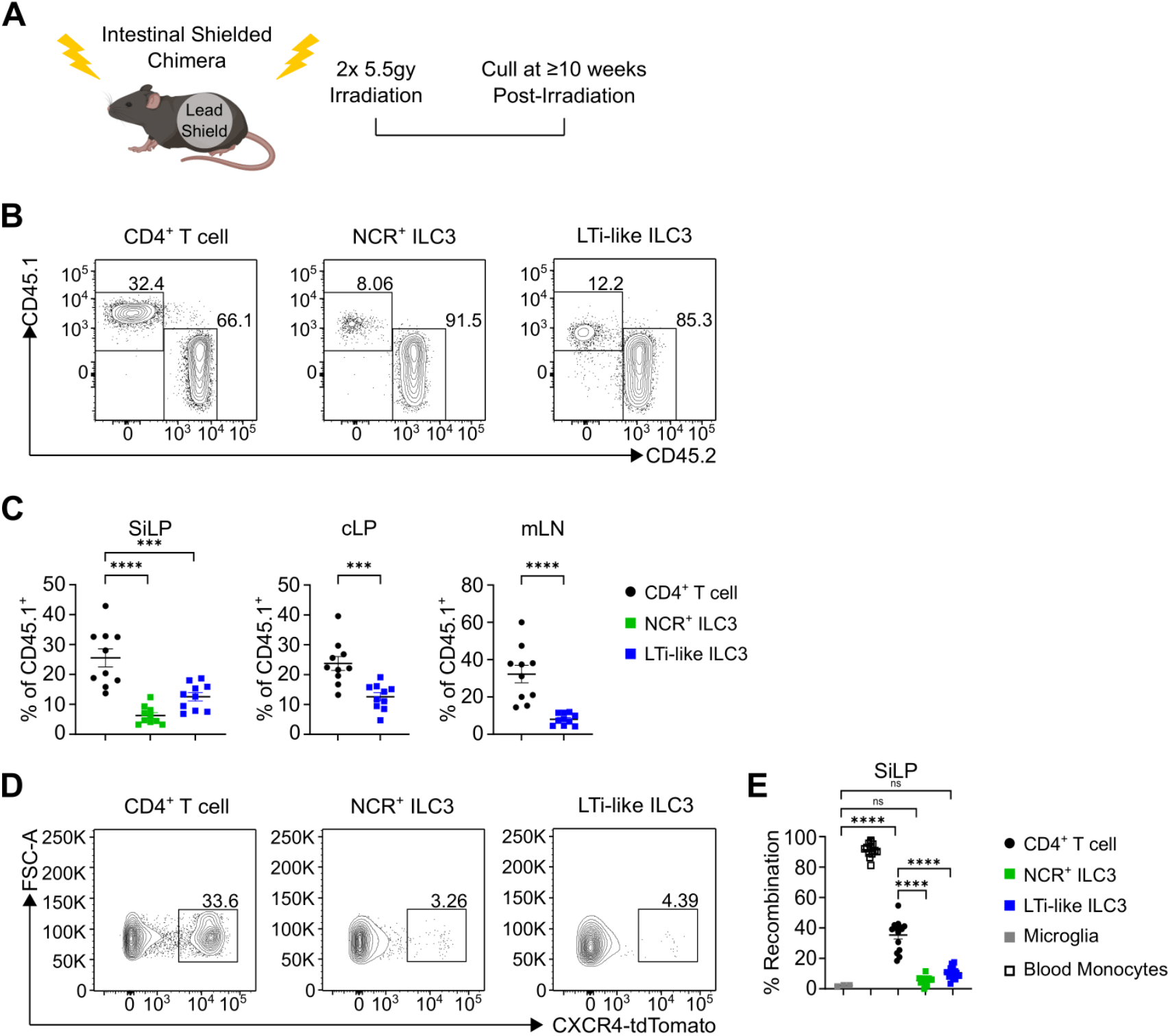
Intestinal ILC3s are maintained independently of bone marrow replenishment. **(A)** Experimental design for intestinal-shielded bone marrow chimeras. Recipient (CD45.2^+^) adult C57BL/6 mice received sub-lethal whole-body irradiation with a lead shield covering the intestines followed by injection of donor (CD45.1^+^) bone marrow. Mice were culled ≥ 10 weeks post irradiation. **(B)** Representative FACS plots of CD45.1^+^ and CD45.2^+^ CD4^+^ T cells (CD3^+^,CD5^+^ / CD4^+^), NCR^+^ ILC3 (Lin^−^, CD127^+^ / RORγt^+^, / NKp46^+^) and LTi-like ILC3 (Lin^−^, CD127^+^/ RORγt^+^ / CCR6^+^) in SiLP at ≥ 10 weeks post-irradiation. **(C)** Frequency of CD45.1^+^ CD4^+^ T cells, NCR^+^ ILC3 and LTi-like ILC3 in SiLP pooled from 3 independent experiments at >10 weeks post-irradiation. **(D** and **E)** Frequency of Cxcr4-^CreERT2-tdTomato^ labelled blood monocytes (CD11b^+^, Ly6C^+^), microglia (CD11b^+^, CD64^+^, F4/80^+^), CD4^+^ T cells, NCR^+^ ILC3s (Lin^−^, CD127^+^, CD90.2^+^/ KLRG-1^−^/ CCR6^−^ NKp46^+^) and LTi-like ILC3s (Lin^−^, CD127^+^, CD90.2^+^/ KLRG-1^−^/ CCR6^+^, NKp46^−^) 12 weeks post-labelling (Microglia, CD4^+^ T cell and ILC3 are normalised to blood monocytes). Data shown as mean +/− SEM and represents two (Microglia in E) or three independent experiments (n = 10 – 14). Numbers in flow plot indicate percentage of cells in the respective gate. *P< 0.05, **P<0.01, ***P<0.001, ****P<0.0001 using unpaired t-test or Tukey’s (C) or Holm-Šídák’s (E) multiple comparisons test.

To circumvent possible confounding factors associated with exposing mice to irradiation, we further tested the contribution of adult bone marrow haematopoiesis to ILC3 replenishment by using a tamoxifen-inducible *Cxcr4*^ERT2Cre-tdTomato^ fate mapping model which induces tdTomato expression in haematopoietic stem cells (HSC) upon tamoxifen administration, and subsequently marks arising progeny [28]. Adult mice were tamoxifen treated and culled 12 weeks post-labelling, and in line with shielded chimera data revealed a clear tdTomato^+^ population to be seen amongst SiLP CD4^+^ T cells (approximately 30%). In contrast, ILC3 subsets had only negligible frequencies of tdTomato^+^ cells (<10%), a rate of recombination that was comparable with other long-lived tissue resident cell populations, including microglia (Figure 1D–1E). Moreover, as expected, blood monocytes exhibited near complete tdTomato labelling-in line with the known rapid turnover and bone marrow replenishment of this population (Figure 1E). Overall, these data suggest mature intestinal ILC3s are subject to minimal replenishment from the bone marrow during adulthood and are long-lived tissue resident cells.

### LTi-like ILC3 are non-proliferative quiescent-like effector lymphocytes

The relatively low contribution of bone marrow input to homeostatic ILC3 maintenance provoked the question as to how tissue-resident ILC3s are maintained in the adult intestine. One way through which immune cell populations may be replenished within tissues is via proliferative self-renewal. Thus, we next determined the proliferative status of intestinal ILC3s at homeostasis by measuring the expression levels of the marker Ki-67. Strikingly, we noticed a clear difference in the frequency of Ki-67 expressing cells between ILC3 subsets. A significant proportion of NCR^+^ ILC3s exhibited Ki-67 expression (Figure 2A-B). Similarly, we observed steady state levels of proliferation in intestinal ILC2 and CD4^+^ T cells (Figure S2A–D). Surprisingly, however, LTi-like ILC3s had only negligible to undetectable expression of Ki-67 (<2%). To further assess the proliferative status of ILC3 subsets, we used a cell cycle dye (Figure 2C, Figure S2E). CD4^+^ T cells and NCR^+^ ILC3 exhibited elevated cell cycle dye staining indicative of cells in the S phase and the G2/M phase stages of the cell cycle, reflecting cell cycle progression towards mitosis (Figure 2D–2E, Figure S2E). Comparatively, LTi-like ILC3 had uniformly low staining for the dye (Figure 2D-E), suggesting very little progression through cell cycle or cell division, reinforcing the idea that LTi-like ILC3 are largely non-proliferative phenotype at steady state. Thus, together our findings suggest LTi-like ILC3 are a quiescent-like cell population under homeostatic conditions.

**Figure 2.**
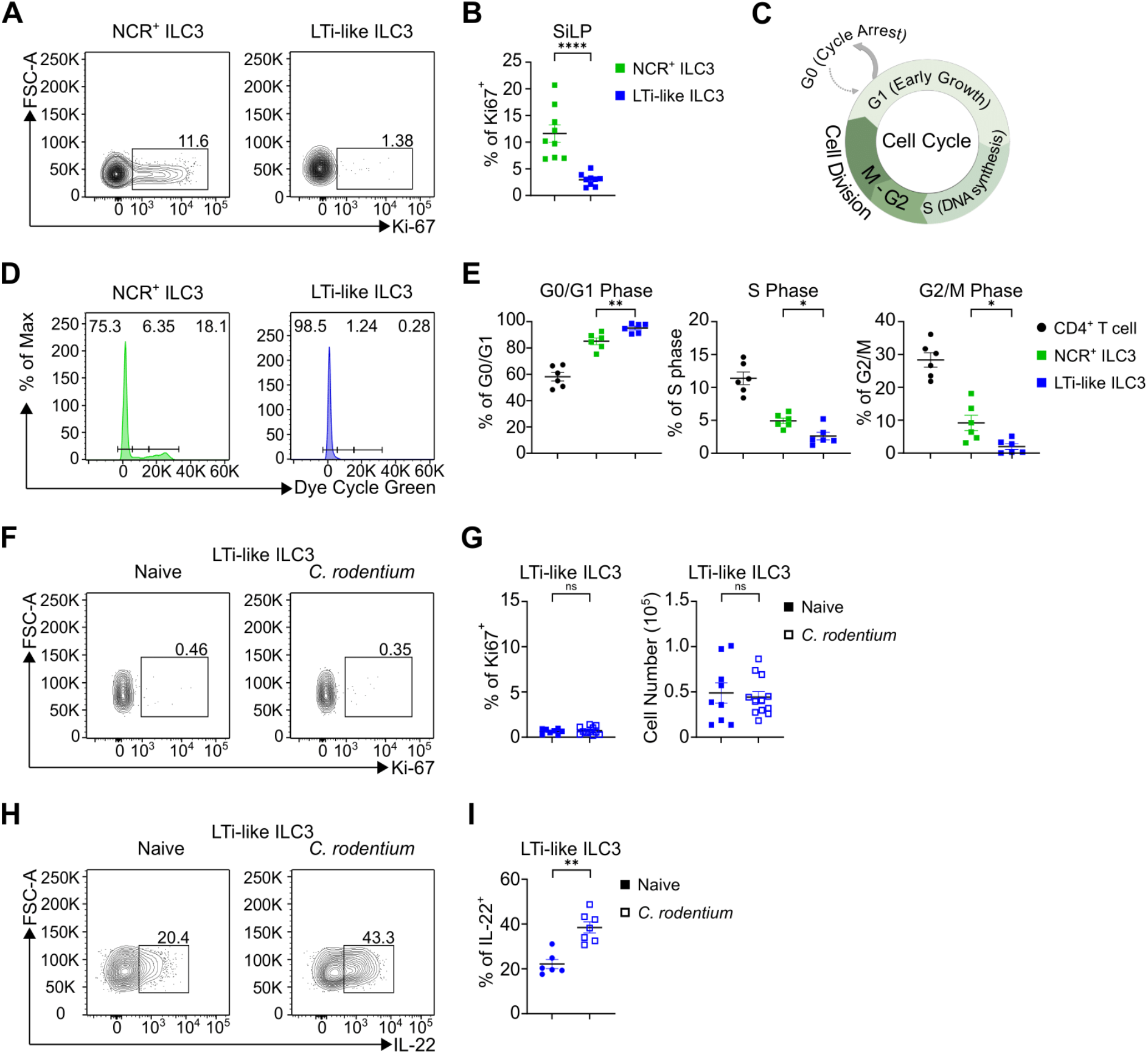
Distinct proliferative capacity of intestinal ILC3 subsets. **(A** and **B)** Frequency of Ki-67 expression in NCR^+^ ILC3s and LTi-like ILC3s in SiLP. **(C)** Cartoon outlining the key cell cycle stages from cycle arrest to mitosis. **(D** and **E)** Frequency of G0/G1 phase, S phase and G2/M phase of CD4^+^ T cells, NCR^+^ ILC3s and LTi-like ILC3s in SiLP using a cell cycle dye. **(F** and **G)** Frequency of Ki-67 expression and cell number in LTi-like ILC3s in cLP of naïve or C. rodentium infected (Day 6) mice. **(H** and **I)** Frequency of IL-22 production in LTi-like ILC3s in cLP of naïve or C. rodentium infected (Day 6) mice. Data shown as mean +/− SEM and represent two (D) or three (B, G, I) independent experiments (n = 6 – 11). Numbers in flow plot indicate percentage of cells in the respective gate. *P< 0.05, **P<0.01, ***P<0.001, ****P<0.0001 using unpaired t-test (B, G) or Mann Whitney test (E, I).

Next, we asked whether LTi-like ILC3 engage proliferation to maintain or expand cell numbers following physiological activation. To assess this we infected mice with the enteric pathogen *C. rodentium* which has been extensively demonstrated to elicit a potent effector cytokine response from which ILC3, and which specifically requires LTi-like ILC3-derived IL-22 for optimal protective immunity [20,29–31]. As a point of comparison, we also assessed the proliferation of TH17 cells, which are also elicited in response to infection. As expected, we observed a significant increase in the proliferation of colonic T_H_17 cells following *C. rodentium* infection (Figure S2F-H). Intriguingly however, we saw no change in the proliferation of LTi-like ILC3s at Day 6 of *C. rodentium* infection in the colon (Figure 2F-G), despite exhibiting an expected and robust increase in IL-22 production from LTi-like ILC3s in response to infection (Figure 2H–I). In contrast Th17 cells did not yet produce significant amounts of IL-22 at this early time point of infection (Figure S2I). Together these results reveal that LTi-like ILC3 are quiescent-like and remain non-proliferative despite acting as important source of effector cytokine during *C. rodentium* infection.

### Induced proliferation specifically depletes LTi-like ILC3

As LTi-like ILC3s did not proliferate at steady state or in response to *C. rodentium* infection, we next sought to understand whether LTi-like ILC3 possessed the capacity to proliferate if exposed to a supra-physiological stimulus. LTi-like ILC3 express high levels of the IL-2Ra subunit CD25 and can outcompete other tissue-resident lymphocytes for available IL-2 [32]. Thus, we treated mice *in vivo* with IL-2 complex (IL-2C), which has previously been shown to promote high levels of proliferation in skin ILCs [33]. Following IL-2C treatment we saw a striking increase in the proliferation of ILC2 and NCR^+^ ILC3 as expected, but also of LTi-like ILC3 - indicating that resident LTi-like ILC3 are capable of proliferation in response to significant stimulation (Figure 3A–D, Figure S3A-C). Increased proliferation correlated with increases in NCR^+^ ILC3 and ILC2 cell frequency as expected, however surprisingly we consistently saw a decrease in LTi-like ILC3 both as a frequency of ILC3 and in total cell numbers following IL-2C treatment, which further coincided with a significant uptake of dead cell dye as well (Figure 3E–I, Figure S3A). Thus, together these data provoke the hypothesis that in contrast to other ILC subsets LTi-like ILC3 may maintain a quiescent state to facilitate survival of in the intestinal tract.

**Figure 3.**
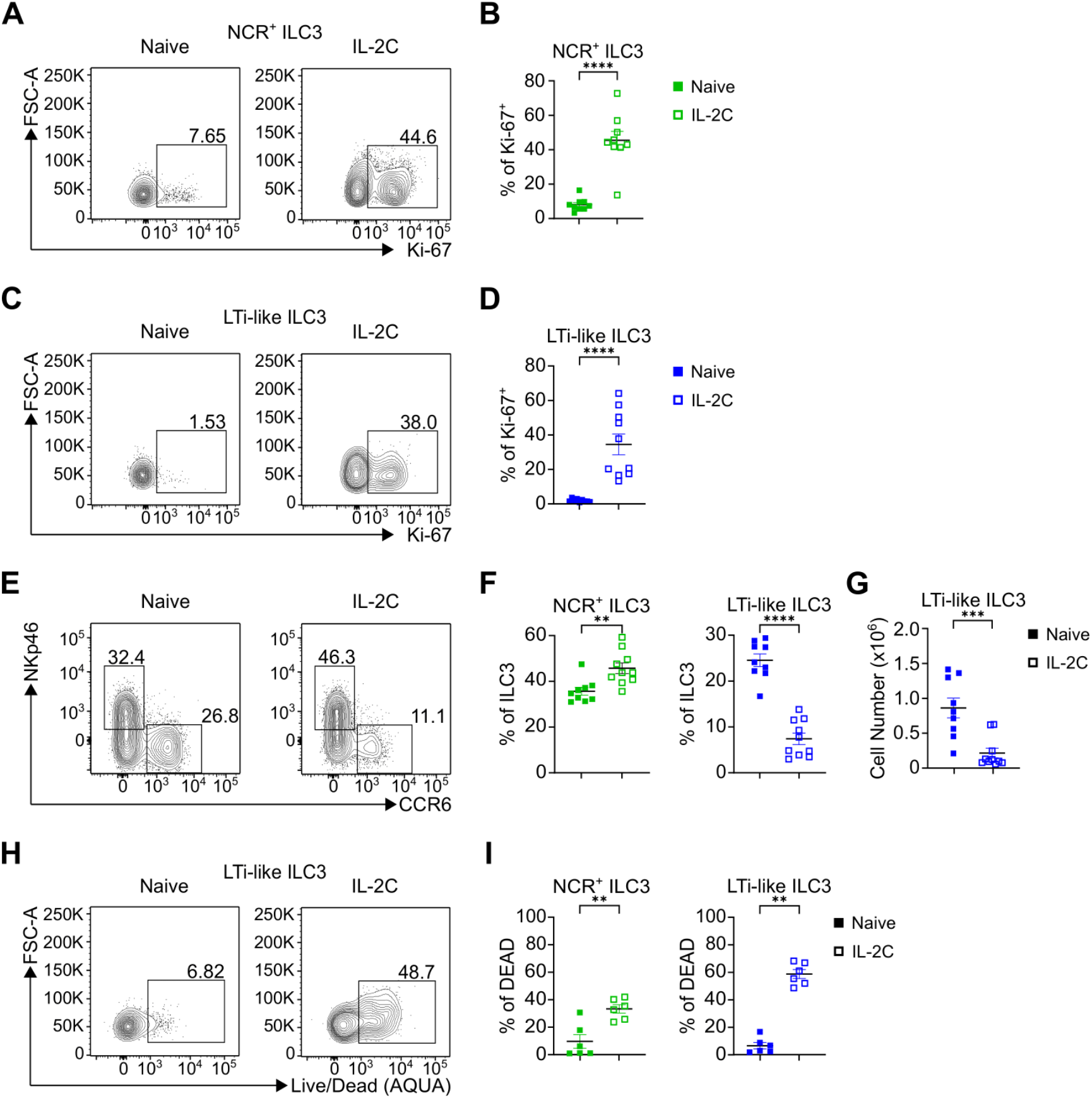
IL-2C induced proliferation results in depletion of LTi-like ILC3. **(A-D)** Frequency of Ki-67 expression in NCR^+^ ILC3 and LTi-like ILC3 in SiLP in naïve or IL-2C treated mice. **(E and F)** Frequency of NCR^+^ ILC3 and LTi-like ILC3 in SiLP in naïve or IL-2C treated mice. **(G)** Cell number of LTi-like ILC3 in SiLP in naïve or IL-2C treated mice. **(H and I)** Frequency of dead NCR^+^ ILC3 and LTi-like ILC3 in SiLP in naïve or IL-2C treated mice. Data shown as mean +/− SEM and represent three independent experiments (n = 6 – 10). Numbers in flow plot indicate percentage of cells in the respective gate. *P< 0.05, **P<0.01, ***P<0.001, ****P<0.0001 using Mann-Whitney test.

### Bcl-2 supports LTi-like ILC3 survival

To determine the underpinning mechanisms through which quiescent LTi-like ILC3 are maintained within tissues for long-term survival, we first generated bulk RNA-Seq data of wild-type siLP NCR^+^ ILC3s and LTi-like ILC3s to identify candidate pathways, and noticed broadly differential relative expression of pro-survival and anti-apoptotic factor gene families between the two ILC3 subsets (Figure 4A). Specifically, LTi-like ILC3 were associated with higher expression of anti-apoptotic genes of the *Bcl2* family when compared to NCR^+^ ILC3, including *Bcl2*, *Mcl1* and *Bcl-w* (Figure 4A). In line with this signature, we could confirm higher expression of Bcl-2 protein on LTi-like ILC3 compared to NCR^+^ ILC3s via flow cytometry (Figure 4B-C). As we had previously observed that IL-2C-driven induction of proliferation led to cell loss, we next determined whether this was associated with changes in Bcl-2 expression. Strikingly, we observed LTi-like ILC3 expressing Ki-67 exhibited a significantly reduced expression of Bcl-2 following IL-2C treatment, further suggesting Bcl-2 is associated with a non-proliferative state in LTi-like ILC3 (Figure 4D-E). To definitively test the importance of Bcl-2 expression in LTi-like ILC3 survival we performed an *ex vivo* culture of SiLP cells with the selective Bcl-2 inhibitor ABT-199. Within 4 hours of incubation with ABT-199 we observed a clear and selective reduction in LTi-like ILC3 in both frequency and cell number, whilst in contrast NCR^+^ ILC3 survival was unaffected (Figure 4F-H), indicating that Bcl-2 activity may act to support LTi-like ILC3 survival. To determine whether inhibition of Bcl-2 was inducing cell death of LTi-like ILC3, we measured the expression of active caspase 3 and uptake of a fixable viability dye. Consistent with the reduction in LTi-like ILC3, we saw elevated expression of active caspase 3 and significantly greater acquisition of dead cell dye within LTi-like ILC3 compared to NCR^+^ ILC3s, confirming that Bcl-2 expression is critical for LTi-like ILC3 survival (Figure 4I–L, Figure S4A-D). Together, these findings identify Bcl-2 expression as a key pro-survival molecule for quiescent LTi-like ILC3s.

**Figure 4.**
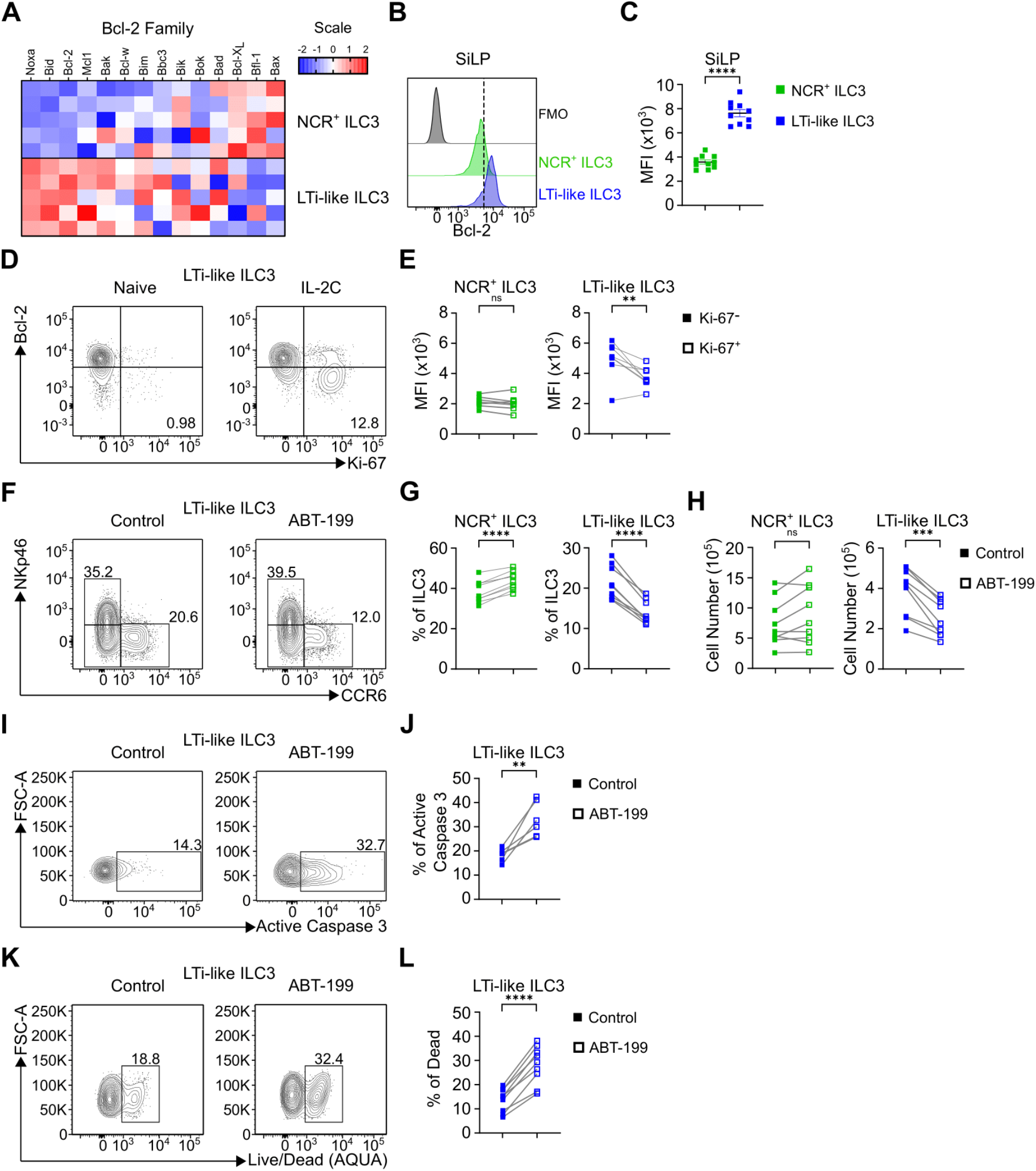
Bcl-2 is a key regulator of LTi-like ILC3 survival. **(A)** Heatmap showing the relative expression levels of Bcl-2 family members between NCR^+^ ILC3s and LTi-like ILC3s from SiLP. **(B-C)** Bcl-2 expression and Bcl-2 MFI of NCR^+^ ILC3s and LTi-like ILC3s in SiLP. **(D)** Bcl-2 and Ki-67 expression in LTi-like ILC3s in SiLP from naïve or IL-2C treated mice. **(E)** Bcl-2 MFI in Ki-67^−^ and Ki-67^+^ populations of NCR^+^ ILC3s and LTi-like ILC3s in SiLP from IL-2C treated mice. **(F**-**H)** Frequency and cell counts of NCR^+^ ILC3s and LTi-like ILC3s in SiLP +/− ABT-199 treatment. **(I-L)** Frequency of active caspase 3 expression (I-J) and live/dead dye acquisition (K-L) in LTi-like ILC3s in SiLP +/− ABT-199 treatment. RNA-Seq data shown as Z-scores relative to overall average. Data shown as mean +/− SEM (C) or individual data points (E, G, H, J, L) and represent two (E and J) or three independent experiments (C, G, H and L) (n = 5 – 10). MFI = mean fluorescence intensity. Numbers in flow plot indicate percentage of cells in the respective gate. *P< 0.05, **P<0.01, ***P<0.001, ****P<0.0001 using unpaired t-test (C) or paired t-test (E, G-H, J, L).

### Bcl-2 protects against OXPHOS-mediated metabolic stress in LTi-like ILC3

Our data suggest that LTi-like ILC3 readily perform effector functions whilst maintaining a quiescent-like state. These findings raise the question as to how the demands of effector function and long-term persistence are met without cellular proliferation and population expansion. Recent studies have highlighted a key role for metabolism in supporting LTi-like ILC3 effector function [34–36]. Thus, we first analysed mitochondrial content and activity and noted that LTi-like ILC3 had comparable levels of mitochondrial mass to analogous Th17 cells (Figure 5A-B). In contrast, LTi-like ILC3 exhibited significantly greater levels of mitochondrial potential (Figure 5C-D), suggesting that LTi-like ILC3 have elevated steady state mitochondrial activity. To determine whether mitochondrial metabolism supported effector function in LTi-like ILC3, we cultured cells with inhibitors of glycolysis and oxidative phosphorylation (OXPHOS). Notably, oligomycin treatment but not 2-DG treatment resulted in considerable ablation of IL-22 production by LTi-like ILC3 (Figure 5E-F), indicating that functionally OXPHOS is utilised by LTi-like ILC3 to elicit effector functions. We next hypothesised that stress associated with the metabolic activity required to fuel IL-22 production may be in part counter-regulated by pro-survival factors. Indeed Bcl-2 family members can mitigate mitochondrial metabolic stress and the production of reactive oxygen species (ROS) to prevent apoptosis [37]. In line with this we detected cellular ROS in LTi-like ILC3 that could be reduced following incubation with oligomycin, but not 2-deoxyglucose (2-DG) (Figure 5G-H; Figure S5A). To determine whether interactions between metabolic stress in part explained the requirement for Bcl-2 in determining LTi-like ILC3 persistence, we cultured LTi-like ILC3 with ABT-199 in the presence or absence of oligomycin. Notably, inhibition of mitochondrial respiration led to a partial rescue of ABT-199 induced cell death in LTi-like ILC3 (Figure 5I-J; Figure S5B), suggesting that metabolic stress from mitochondrial respiration required to maintain effector functions, is in part negated by expression of Bcl-2 to prevent apoptosis (Figure S5C).

**Figure 5.**
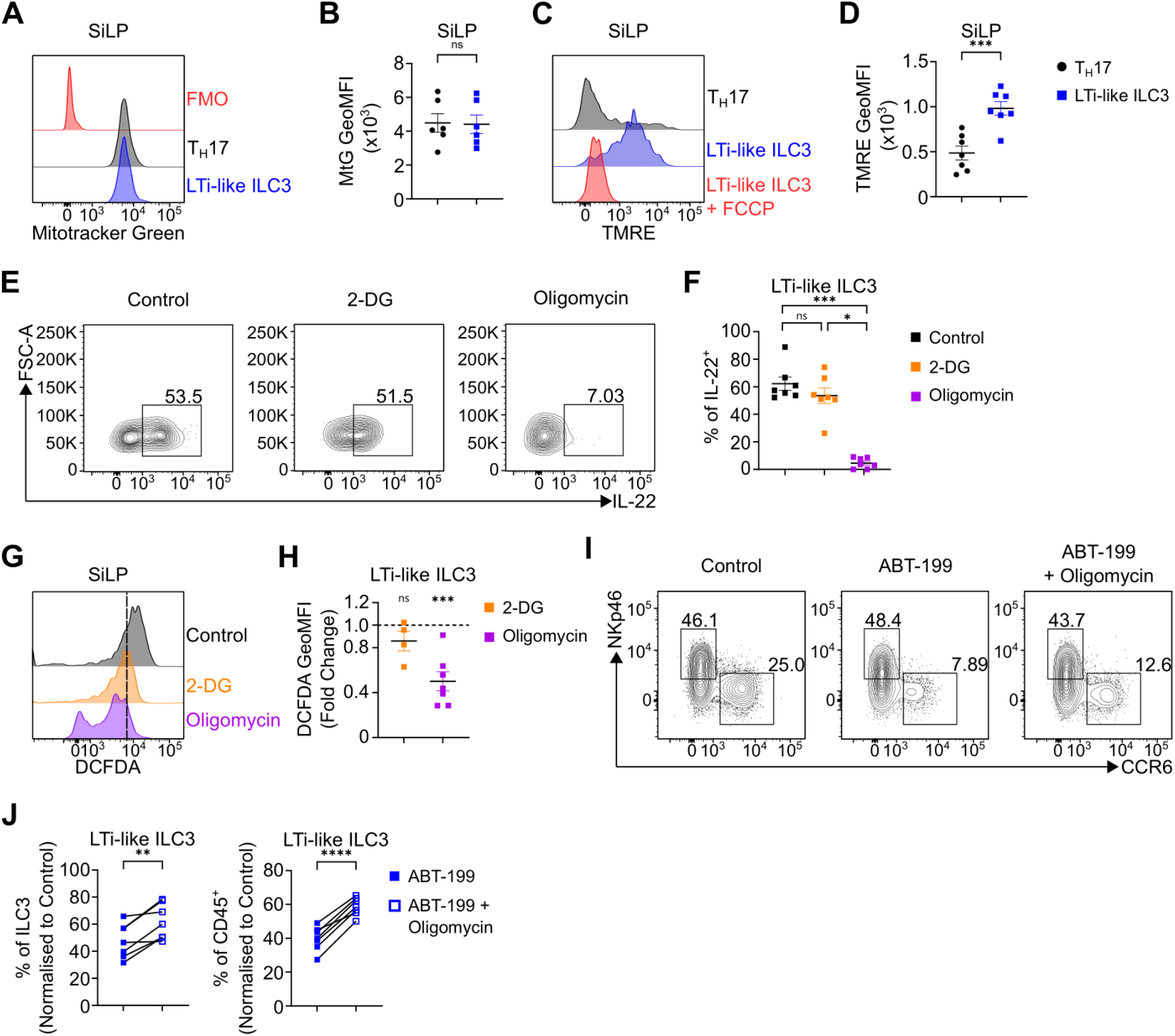
Bcl-2 protects against OXPHOS-mediated metabolic stress. **(A-B)** Representative histogram of mitochondrial mass and MFI in TH17 cells and LTi-like ILC3 in SiLP. **(C-D)** Representative histogram of mitochondrial activity/polarisation and MFI in TH17 cells and LTi-like ILC3 in SiLP. **(E-F)** IL-22 production +/− 2-DG or Oligomycin treatment in LTi-like ILC3 in SiLP. **(G-H)** Representative histogram of cellular ROS (DCFDA) and DCFDA MFI fold change relative to control following 2-DG or Oligomycin treatment in LTi-like ILC3 in SiLP. **(I-J)** Frequency of LTi-like ILC3 +/− ABT-199 and Oligomycin treatment in SiLP. Data shown as mean +/− SEM (B, D, F, H) or individual data points (J) and represent two (F, 2-DG in H) or three independent experiments (B, D, H, J) (n = 4 – 7). MFI = geometric mean fluorescence intensity. Numbers in flow plot indicate percentage of cells in the respective gate. *P< 0.05, **P<0.01, ***P<0.001, ****P<0.0001 using unpaired t-test (B, D, F), paired t-test (J) or Dunn’s multiple comparisons test (H).

## Discussion

Here, we show that LTi-like ILC3 in the gastrointestinal tract are maintained largely independent of hematopoietic replenishment at steady state through a pro-survival program that negates the stress of effector function in otherwise quiescent cells. Moreover, our data further highlight the differential biology between LTi-like ILC3 and other ILC subsets, including NCR^+^ ILC3. Strikingly, we demonstrate that LTi-like ILC3 are typically non-proliferative cells, both at steady state and during a robust effector cytokine response to *C. rodentium* infection, and instead are associated with high levels of the pro-survival molecule Bcl-2 that protects these cells against metabolic stress.

The ontogeny of LTi-like ILC3 has until recently been unclear. Fetal LTi that arise during embryogenesis were previously shown to persist, but also to be partially replaced immediately after birth [12]. However, whether hematopoietic input continues to replenish ILC3 subsets once populations have been fully established in adulthood remained unclear. He were demonstrate relatively low levels of replenishment of LTi-like ILC3 during adulthood and persistence of mature tissue-resident ILC3 for periods of 10 weeks and over, in contrast with early reports estimating the half-life of LTi-like ILC3 at 22-26 days [21]. One possible explanation for the discrepancy between these estimates could be the life-stage in which ILC3 turnover was determined. For example, Sawa *et al* determined turnover of ILC3 during the initial neonatal period of seeding and weaning, whereas here we focused only on established ILC3 populations in the adult intestinal tissue. Indeed, as discussed above a recent key study suggested that while many ILC3 are seeded prior to birth a further wave of haematopoiesis may contribute to the ILC3 pool in these first weeks of life [12]. Adding to these prior studies, our findings suggest that while haematopoiesis may contribute to the initial colonization of intestinal tissues by ILC3 subsets, this may be limited to early life - with only negligible turnover and hematopoietic contribution beyond weaning and in the adult gut.

Our data further suggest LTi-like ILC3 are a quiescent-like population that are unlikely to be maintained through homeostatic proliferation and self-renewal. Surprisingly, LTi-like ILC3 were found to be refractory to proliferation even during infection with *C. rodentium*. Moreover, although IL-2C treatment could induce proliferation of LTi-like ILC3 it also surprisingly led to a specific loss of LTi-like ILC3. The reasons for this latter observation are not yet fully clear, however the correlation with Bcl-2 downregulation could in part explain why proliferation leads to loss of LTi-like ILC3 *in vivo*. This biology is reminiscent of previous findings demonstrating high expression of Bcl-2 in non-cycling NK cells which is subsequently lost upon entry into cell cycle [38]. Moreover, the identification of a strong pro-survival program in LTi-like ILC3 characterised by high levels of Bcl-2 is in line with previous observations [39], and intriguingly similar to that described for other quiescent intestinal immune cell populations such as tissue-resident memory T cells [40].

The transcriptional or environmental cues that imprint and maintain a Bcl-2-associated and quiescent-like phenotype on LTi-like ILC3 remain unclear. However, prior studies have demonstrated that inflammatory cues or genetic-induced deletion of niche-retaining chemokine receptor expression drive the mobilisation of LTi-like ILC3 away from organised lymphoid structures, such as cryptopatches and isolated lymphoid follicle niches in the gut, and into the lamina propria where they become proliferative [17,41,42]. These findings suggest tissue niche may be critical for the phenotype, long-term survival and persistence of LTi-like ILC3. Indeed, unlike NCR^+^ ILC3, LTi-like ILC3 are typically restricted to lymphoid structures in healthy animals where they co-localize with stromal cells that provide survival cues such as IL-7 and stem cell factor [43]. Therefore, future studies are required to determine if quiescence and persistence of LTi-like ILC3 in the gut are directly dependent upon the cellular niche and tissue microenvironment in which they are localized.

Together our findings suggest LTi-like ILC3 are a quiescent long-lived cell with minimal turnover at steady-state. Prior studies demonstrated that an intestinal resident RORγt^+^ T-bet^+^ cell population lacking expression of mature ILC3 subset markers acts as an immature pool of precursors for NCR^+^ ILC3, but demonstrated no capacity to transdifferentiate into LTi-like ILC3 [24]. These findings further emphasise the developmental, phenotypic and functional distinction between these two ILC3 subsets [15], and are in line with evidence to demonstrate the earlier divergence of LTi precursors during development [44], and lack of LTi-like ILC3 progeny generated from the common helper ILC progenitor – which is conversely able to give rise to NCR^+^ ILC3 [45]. Thus, while we cannot completely rule out the contribution of an as-yet unidentified tissue-resident progenitor to LTi-like ILC3 maintenance, our data together with current understanding would suggest cell-intrinsic survival mechanisms favour long-term persistence of LTi-like ILC3 without significant input from peripheral or tissue-resident progenitors.

Finally, we highlight ILC-intrinsic metabolism as a critical determinant of effector function and cell fate, in line with a number of recent studies [34,35,46]. Here we demonstrate mitochondrial respiration is required to support effector cytokine production and that, in the absence of Bcl-2-mediated protection, metabolic activity may lead to cell death. In contrast, IL-2 driven proliferation resulted in a reduction in LTi-like ILC3 and intrinsic Bcl-2 expression. A previous study demonstrated IL-2 induction of ILC3 proliferation requires mTOR [36], suggesting such signals may remodel the metabolic program of ILC3 to facilitate these demands [35]. One possibility is that the metabolic stress associated with proliferation may initially be detrimental to LTi-like ILC3 survival, but that cells which persist in this initial stimulation may undergo significant metabolic reprogramming. Indeed, one recent report suggested metabolic training may reprogram ILC3 to become more proliferative in response to repeat challenges [34].

Taken together our data further emphasise the importance of considering ILC3 subset specific biology and advance our understanding the molecular mechanisms that underpin their survival, persistence and effector functions. Further studies to uncover the interplay between tissue-niche, cell-intrinsic metabolism and ILC3 subset dynamics will help to uncover the contributions of these cells to tissue health or chronic inflammatory diseases in the gastrointestinal tract.

## Materials and Methods

### Mice

Age- and sex-matched mice between 8 to 12 weeks of age were used. *Cxcr4*^CreERT2^ ROSA^tdTomato^ (ROSA^tdT^) [28], *Id2*^CreERT2^ ROSA^tdRFP^, [26,27] mouse models have been previously described. *Cxcr4*^CreERT2^ ROSA^tdT^ were maintained at the Queens Medical Research Institute (Edinburgh, UK), while all other strains were housed at the University of Manchester (Manchester, UK). All mice were on the C57BL/6 background and were maintained under specific pathogen-free conditions, and provided with food and water ad libitum. C57BL/6 (Envigo) female mice were ordered from Jackson Laboratory.

### Tamoxifen treatment

ID2-^CreERT2^ ROSA^tdRFP^ reporter model, 8-week-old mice received 5 doses of 5mg tamoxifen diluted in 5μl of ethanol and 95μl of corn oil via oral gavage on Days 0, 2, 4, 7 and 9. Mice were culled either 10 days or 52 days post-final gavage. CXCR4-CreERT2 ROSA^tdT^ reporter model, 6-8-week-old mice received 3 doses of 0.12mg tamoxifen per gram of body weight for 3 consecutive days in a 100μl bolus. Mice were culled 12 weeks post-final gavage.

### IL-2 Complex treatment

IL-2 complex (IL-2C) was made by mixing 2.5ug of IL-2 monoclonal antibody (JES6-1A12, 2B Scientific) with 7.5ug recombinant IL-2 (ThermoFisher) in 200ul of PBS for 20 minutes at 37 °C. 3 doses of IL-2C were injected via i.p over 6 days and mice were sacrificed 2 days after the last dose.

### Bone marrow chimeras

For intestinal shielded bone marrow chimera, CD45.2^+^ C57BL/6 mice (Envigo laboratories), were selectively irradiated by placing a lead shield over the abdomen and irradiating the body with 2 doses of 5.5 grays of radiation, 5 minutes per dose. Following irradiation, mice were reconstituted with CD45.1^+^ congenic bone marrow via i.v. and mice were culled and tissues harvested as indicated. For whole body bone marrow chimera, CD45.2^+^ C57BL/6 mice (Envigo laboratories) were irradiated using the same approach without the lead shield. Mice were then reconstituted with 50/50 CD45.1^+^/CD45.2^+^ bone marrow via i.v. Mice were culled 32 weeks later.

### *C. rodentium* culture and infection

*C. rodentium* (Nalidixic acid resistant strain) was originally a kind gift from Gad Frankel (Imperial College London). *C. rodentium* was cultured from a glycerol stock in 10ml of LB Broth containing nalidixic acid (50ug/ml) for 12-18 hours at 37°C, shaking at 200 r.p.m. The resulting bacteria was then streaked onto an LB Agar plate containing nalidixic acid and left at 37 °C for 24 hours. Singles colonies were picked and grown in 10ml of LB Broth containing nalidixic acid for 12-18 hours. Bacteria were then centrifuged at 3000xg for 10 minutes and resuspended in 1ml of sterile PBS. Infective doses (~ 2×10^9^ CFU) of *C. rodentium* were confirmed via plating and administered to mice via oral gavage.

### Tissue digestion

Intestinal lamina propria lymphocytes were isolated by first removing the intestinal tissue from euthanized mice, removing fat and Peyer’s patches and cutting longitudinally. Luminal contents were removed by vortexing in PBS and then the epithelial lining was removed by vortexing in PBS containing 1mM EDTA, 1mM dithothreitol and 5% FBS for 10 minutes, and a subsequent second 15 minute incubation with constant agitation at 37 °C. Tissue was then digested in 10ml of complete media (10% FBS, 100 units/ml penicillin, 100ug/ml streptomycin, 2mM L-glutamine in RMPI media) containing 20ug/ml DNase I (Sigma-Aldrich) and either for Collagenase VIII for 25 minutes, or Collagenase/Dispase for 45 minutes, vortexing at 37 °C. Collagenase Dispase was used for experiments where phenotyping of ILC3 was performed whilst Collagenase VIII was used to increase cell yields for ex vivo assays. Subsequently, digested tissue was filtered through a 100uM nylon filter and a 70uM nylon filter. Filtrate was then centrifuged at 500xg for 5 minutes to pellet lamina propria lymphocytes.

### Flow Cytometry

Single-cell preparations were stained with the following antibodies:

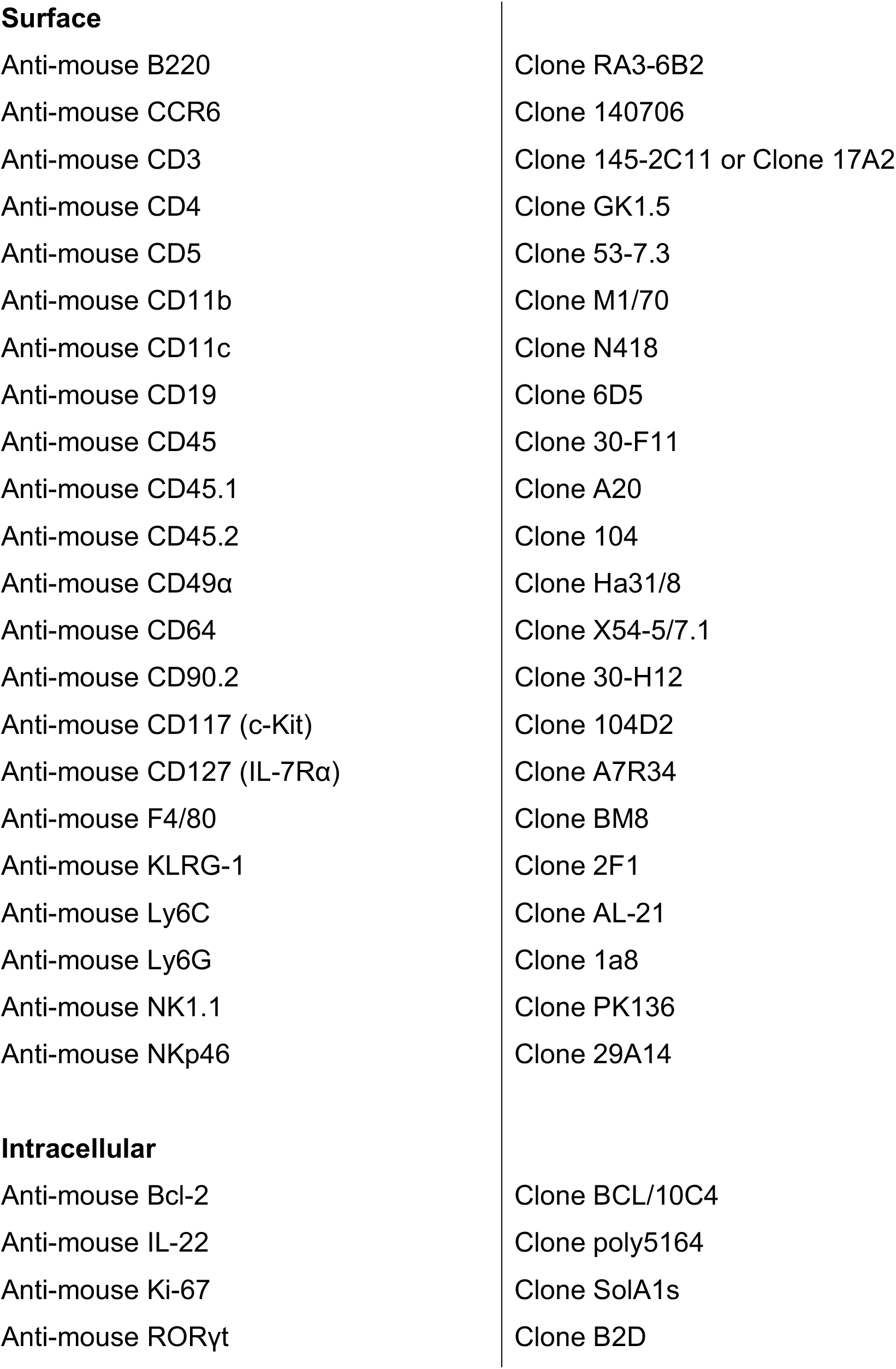

**Surface**
|  |  |
| --- | --- |
| Anti-mouse B220 | Clone RA3-6B2 |
| Anti-mouse CCR6 | Clone 140706 |
| Anti-mouse CD3 | Clone 145-2C11 or Clone 17A2 |
| Anti-mouse CD4 | Clone GK1.5 |
| Anti-mouse CD5 | Clone 53-7.3 |
| Anti-mouse CD11b | Clone M1/70 |
| Anti-mouse CD11c | Clone N418 |
| Anti-mouse CD19 | Clone 6D5 |
| Anti-mouse CD45 | Clone 30-F11 |
| Anti-mouse CD45.1 | Clone A20 |
| Anti-mouse CD45.2 | Clone 104 |
| Anti-mouse CD49α | Clone Ha31/8 |
| Anti-mouse CD64 | Clone X54-5/7.1 |
| Anti-mouse CD90.2 | Clone 30-H12 |
| Anti-mouse CD117 (c-Kit) | Clone 104D2 |
| Anti-mouse CD127 (IL-7Rα) | Clone A7R34 |
| Anti-mouse F4/80 | Clone BM8 |
| Anti-mouse KLRG-1 | Clone 2F1 |
| Anti-mouse Ly6C | Clone AL-21 |
| Anti-mouse Ly6G | Clone 1a8 |
| Anti-mouse NK1.1 | Clone PK136 |
| Anti-mouse NKp46 | Clone 29A14 |
| **Intracellular** |  |
| Anti-mouse Bcl-2 | Clone BCL/10C4 |
| Anti-mouse IL-22 | Clone poly5164 |
| Anti-mouse Ki-67 | Clone SolA1s |
| Anti-mouse RORγt | Clone B2D |

Dead cells were excluded from analysis using the LIVE/DEAD Fixable Aqua Dead Cell Stain Kit (Invitrogen). Lineage exclusion gating to identify ILC was CD3^−^, CD5^−^, NK1.1^−^, B220^−^, CD11b^−^ and CD11c^−^.

Staining of intracellular markers was performed by using the Foxp3 Transcription Factor Buffer kit (eBioscience). For analysis of cell cycle status, live single-cell preparations were stained with 10μM Vybrant™ Dye Cycle™ (ThermoFisher) for 30 minutes at 37°C, 5% CO2. For analysis of mitochondrial mass, live single-cell preparations were stained with 100nM MitoTracker™ Green (ThermoFisher) for 30 minutes in HBSS. For analysis of mitochondrial potential/activity, live single-cell preparations were stained 200nM of TMRE (Abcam) for 30 minutes in HBSS; As a negative control, live single-cell preparations were co-treated with 50μM of carbonyl cyanide 4-(trifluoromethoxy) phenylhydrazone (FCCP) for 30 minutes in HBSS. For analysis of cellular ROS, live single-cell preparations were stained with 20μM 2’,7’ – dichlorofluorescin diacetate (DCFDA, Abcam) for 5 minutes in HBSS; As a positive control live single-cell preparations were co-treated with 200μm of tert-butyl hydrogen peroxide (TBHP) for 5 minutes in HBSS. All flow cytometry data was acquired on a BD Fortessa and analysed using FlowJo version 10.

### *Ex vivo* assays

#### Ex Vivo ILC3 cytokine stimulation

5×10^6^ SiLP cells were incubated in 250ul of complete media containing 20ng/ml IL-1β and IL-23 (R&D Systems) for 2 hours at 37°C, 5% CO_2_. After 2 hours, cells received 50ul of complete media containing eBioscience™ Cell Stimulation Cocktail plus protein transport inhibitors (ThermoFisher) (2μl/ml) on top and incubated for a further 3 hours. To measure the impact of metabolic inhibition on cytokine production, 2-deoxyglucose (2.5mM) or Oligomycin (1μM) was added when adding IL-1β/-23. After incubation, cells were centrifuged, and cells were used for antibody staining for flow cytometry.

#### Bcl-2 inhibitor incubation

5×10^6^ SiLP cells were incubated in 250ul complete media containing 200nM ABT-199 (Stratech) for 4 hours at 37°C, 5% CO2. To measure the impact of metabolic inhibition on Bcl-2 mediated survival, Oligomycin (1uM) was added when adding ABT-199. After incubation, cells were centrifuged, and cells were used for antibody staining for flow cytometry. To assess active caspase 3 activity the CaspGLOW™ Fluorescein Active Caspase-3 Staining Kit (Invitrogen) was used. Cells were incubated in ABT-199 as stated above but incubated with 2.5ul of DEVD-FMK (FITC-conjugated Caspase 3 inhibitor) in the final hour. Cells were then centrifuged, cells were used for antibody staining for flow cytometry.

### RNA-Seq

RNA was isolated from sort-purified NCR^+^ ILC3 and LTi-like ILC3 from SiLP using an RNA Purification Kit (Norgen) and amplified using SMART-Seq™ v4 Ultra Low Input RNA Kit for sequencing (Takara Bio USA, Inc., Mountain View, USA), producing double stranded cDNA. cDNA was purified with AMPure XP beads and quantified with Qubit (Life Technologies™). Library preparation was done using NEBNext Ultra RNA Library Prep Kit (New England Biolabs, Ipswich, USA) following manufacturer recommendations. Libraries were then sequenced on Illumina NovaSeq 6000 S4 flowcell with PE150 according to results from library quality control and expected data volume. CASVA based recognition was used to convert raw counts into FASTQ files. Reads with adapter contamination, >10% uncertain nucleotides or had a base quality of <5 in over 50% of the read were filtered out. Remaining reads were aligned with HISAT2. Differential expression analysis was performed using R software and DeSeq2 package. Z-scores were calculated from the resulting differential expression output.

### Statistics

All graphs and statistical analysis were carried out in GraphPad Prism 7. Group *n* numbers are stated in each figure legend along with statistical test utilized. Parametric or non-parametric tests were based on Shapiro-Wilk normality test outcome. Mean fluorescence intensities were calculated via FlowJo v 10.6.1.

## Acknowledgements

The authors acknowledge members of the Hepworth lab for critical discussion. Gareth Howell, David Chapman and the University of Manchester flow cytometry core for support. We also thank Xavier Romero Ros, Josquin Nys and Suzanne Cohen (AstraZeneca) for financial and scientific support as part of an MRC CASE PhD studentship awarded to James King. Research in the Gentek Laboratory is supported by a Kennedy Trust for Rheumatology Research Senior Fellowship and a Cancer Research UK Immunology Project Award. Research in the Hepworth Laboratory is supported by a Sir Henry Dale Fellowship jointly funded by the Wellcome Trust and the Royal Society (Grant Number 105644/Z/14/Z), a BBSRC responsive mode grant (BB/T014482/1) and a Lister Institute of Preventative Medicine Prize.

## Author contributions

J.I.K – Conceptualization, investigation, data curation, formal analysis, methodology, visualization, writing, reviewing and editing of manuscript.

F.M.G., B.M-D., R.T-P., M.S.M, S.H.H - investigation, data curation, formal analysis.

X.RR. – Resources, Supervision

R.G – Resources, Conceptualization, supervision, Validation.

M.R.H - Conceptualization, investigation, data curation, formal analysis, visualization, funding acquisition, project administration, supervision, writing, reviewing and editing of manuscript.

---

**Figure S1.**
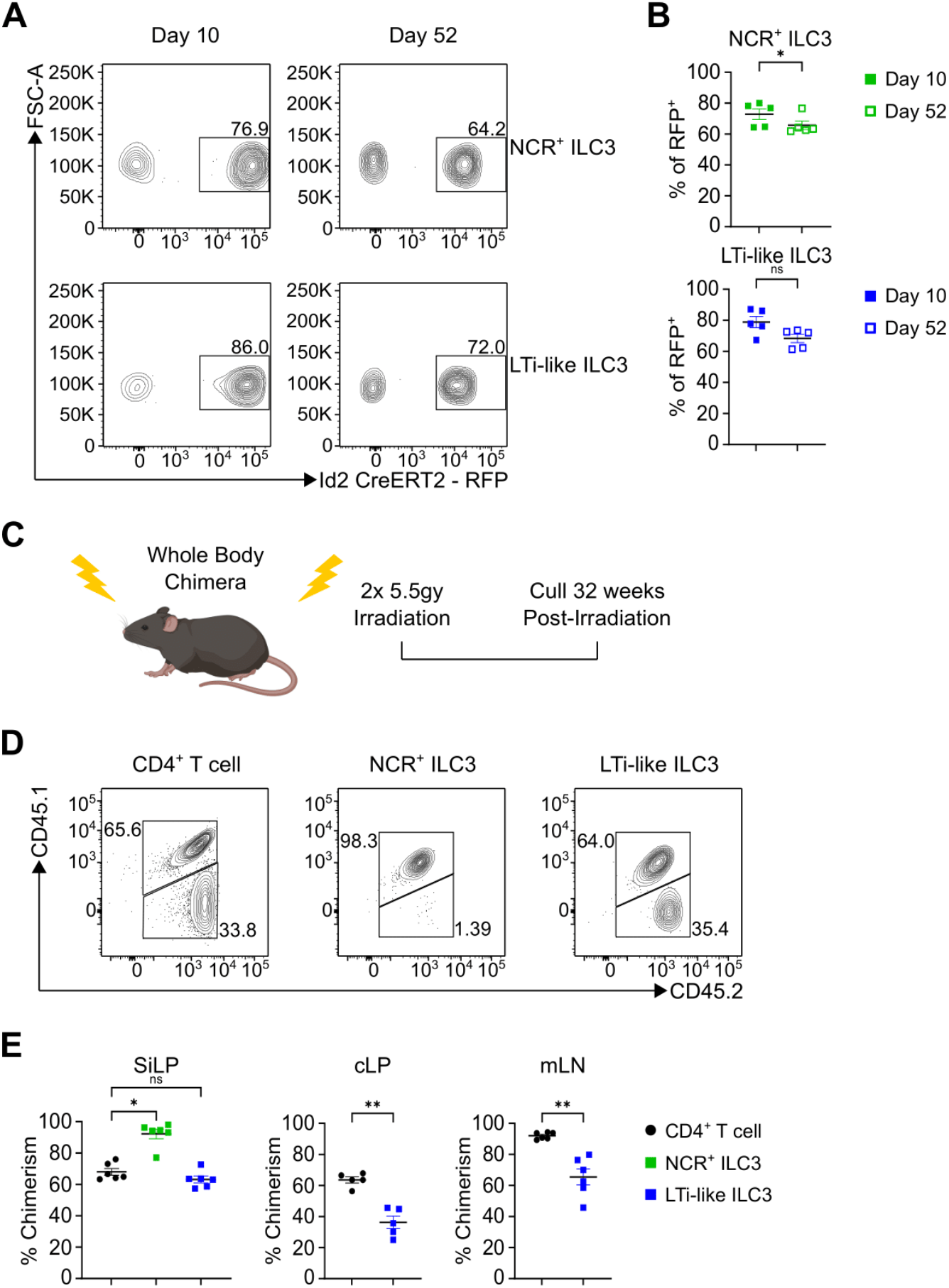
Intestinal ILC3s are maintained independently of bone marrow replenishment. **(A-B)** Frequency of Id2^ERT2Cre-RFP^ labelled NCR^+^ ILC3s and LTi-like ILC3s in SiLP 10 or 52 days post-labelling. **(C)** Experimental design for whole body chimeras. Recipient (CD45.2^+^) adult C57BL/6 mice received sub-lethal whole body irradiation followed by injection of Donor (CD45.1/2^+^) bone marrow. Mice were culled 32 weeks post irradiation. **(D** and **E)** Frequency of CD45.2^+^ CD4^+^ T cells (CD3^+^, CD5^+^/CD4^+^), NCR^+^ ILC3s (Lin^−^, CD127^+^/ RORγt^+^, NKp46^+^) and LTi-like ILC3s (Lin^−^, CD127^+^/ RORγt^+^, CCR6^+^) in SiLP. Data shown as mean +/− SEM and represent two independent experiments (n = 5 – 6). Numbers in flow plot indicate percentage of cells in the respective gate. *P< 0.05, **P<0.01, ***P<0.001, ****P<0.0001 using Dunn’s multiple comparisons test (B) or Mann-Whitney test (D).

**Figure S2.**
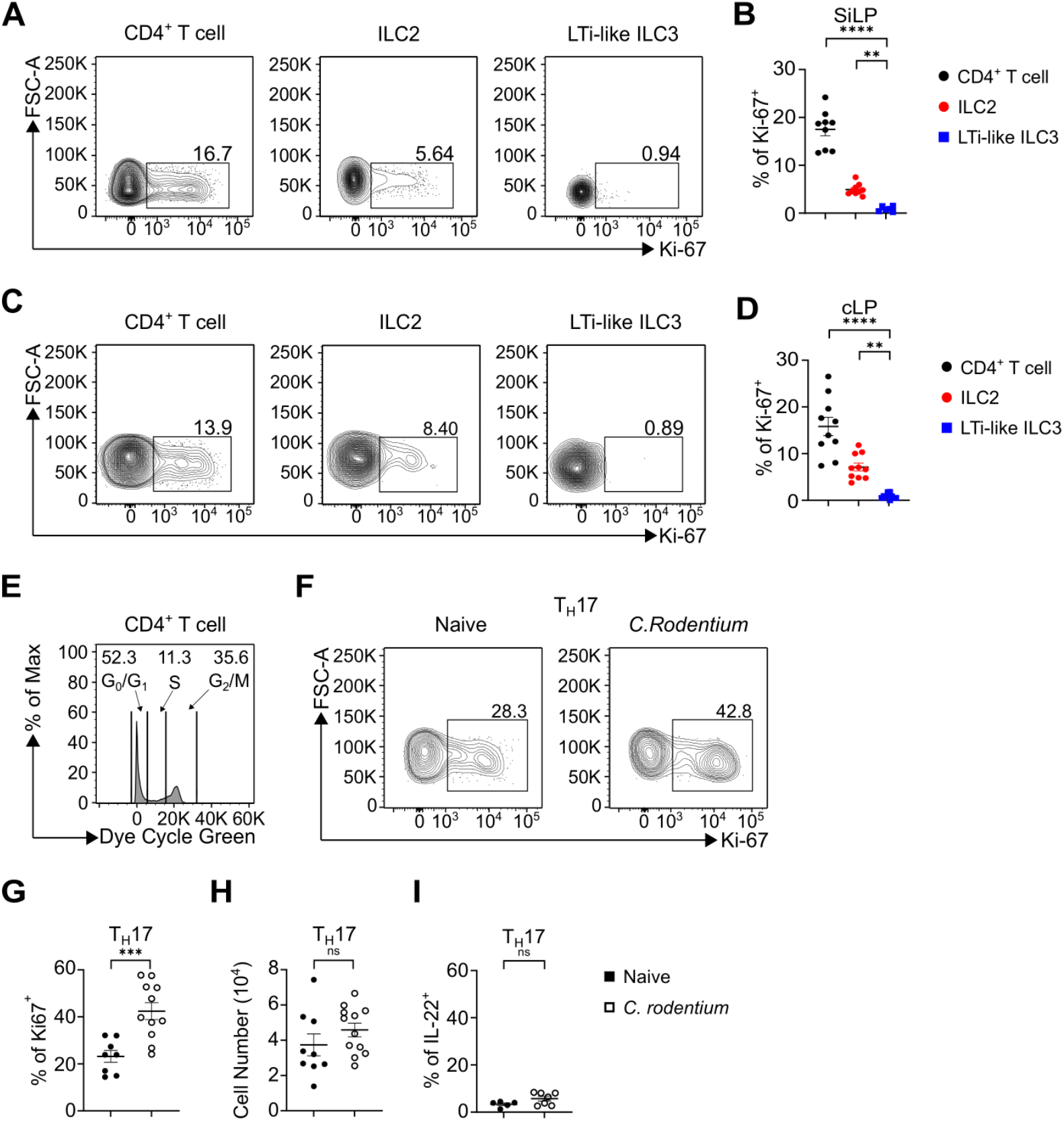
Proliferative capacity of intestinal ILC and T cells. **(A** and **B)** Frequency of Ki-67 expression in CD4^+^ T cell, ILC2 and LTi-like ILC3 in SiLP or **(C** and **D)** cLP at steady state. **(E)** Gating strategy of cell cycle stages using cell cycle dye. **(F and G)** Frequency of Ki-67 expression and cell number **(H)** in TH17 cell in cLP of naïve or C. rodentium infected (Day 6) mice. **(I)** Frequency of IL-22 production in TH17 cell in cLP of naïve or C. rodentium infected (Day 6) mice. Data shown as mean +/− SEM and represent two (B, D) or three (F, G) independent experiments (n = 5 – 12). Numbers in flow plot indicate percentage of cells in the respective gate. *P< 0.05, **P<0.01, ***P<0.001, ****P<0.0001 using Holm-Šídák’s multiple comparisons test (B, D), unpaired t-test (F, G) or Mann-Whitney test (I).

**Figure S3.**
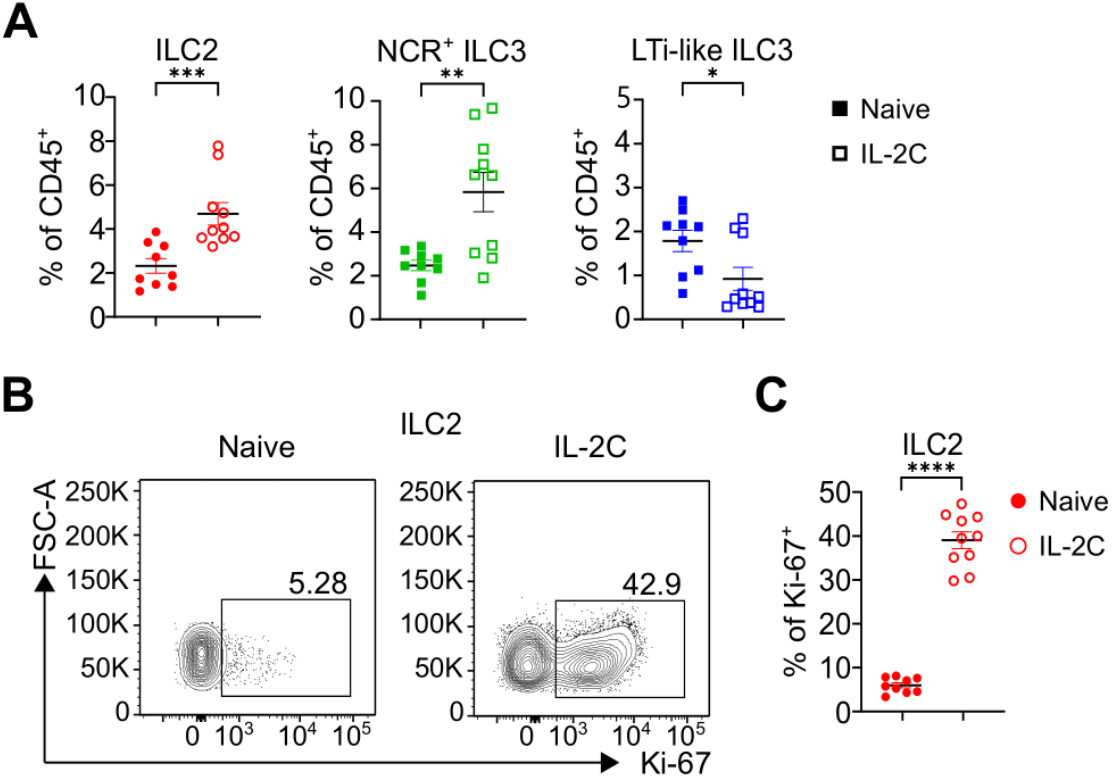
IL-2C-induced proliferation induces proliferation of ILC but increases frequency of cell-death in ILC3. **(A)** Frequency of ILC2, NCR^+^ ILC3 and LTi-like ILC3 in SiLP in naïve or IL-2C treated mice as percentage of CD45^+^ lymphocytes. **(B-C)** Frequency of Ki-67 expression in ILC2 in SiLP in naïve or IL-2C treated mice. Data shown as mean +/− SEM and represent three independent experiments (n = 9 – 10). Numbers in flow plot indicate percentage of cells in the respective gate. *P< 0.05, **P<0.01, ***P<0.001, ****P<0.0001 using Mann-Whitney test.

**Figure S4.**
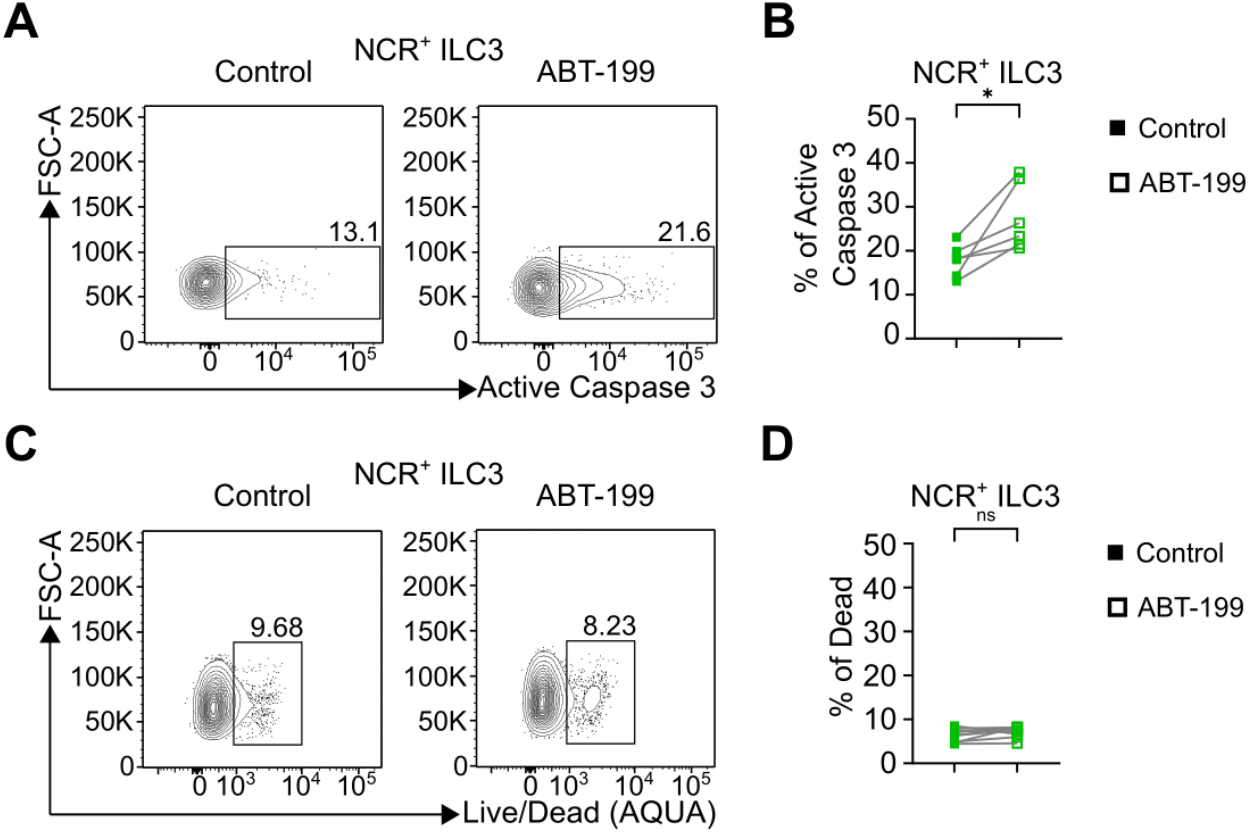
Bcl-2 inhibition has little effect on NCR^+^ ILC3 survival. **(A-D)** Frequency of active caspase 3 expression (A-B) and live/dead dye acquisition (C-D) in NCR^+^ ILC3 in SiLP +/− ABT-199 treatment. Data shown as individual data points and represent two (B) or three independent experiments (D) (n = 5 – 9). Numbers in flow plot indicate percentage of cells in the respective gate. *P< 0.05, **P<0.01, ***P<0.001, ****P<0.0001 using paired t-test.

**Figure S5.**
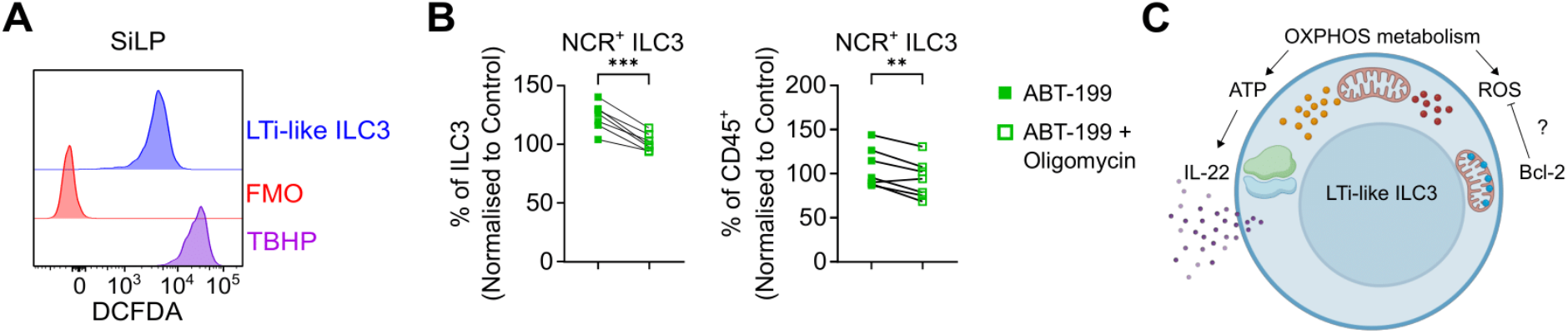
Intracellular balance between metabolism and stress in ILC3. **(A)** Representative histogram of cellular ROS (DCFDA) +/− FMO or TBHP in LTi-like ILC3 in SiLP. **(B)** Frequency of NCR^+^ ILC3 +/− ABT-199 and Oligomycin treatment in SiLP. **(C)** Cartoon detailing the balance of metabolic stress in LTi-like ILC3 to facilitate function but also survival. Data shown as mean +/− SEM and represent three independent experiments) (n = 6). *P< 0.05, **P<0.01, ***P<0.001, ****P<0.0001 using paired t-test.

